# Atypical beta-band effects in children with dyslexia in response to rhythmic audio-visual speech

**DOI:** 10.1101/2023.03.29.534542

**Authors:** Mahmoud Keshavarzi, Kanad Mandke, Annabel Macfarlane, Lyla Parvez, Fiona Gabrielczyk, Angela Wilson, Usha Goswami

**Affiliations:** Centre for Neuroscience in Education, Department of Psychology, University of Cambridge, Cambridge, CB2 3EB, United Kingdom

**Keywords:** Developmental dyslexia, phase entrainment, phase-amplitude coupling, beta band, rhythmic audio-visual speech

## Abstract

Children with dyslexia are known to show impairments in perceiving speech rhythm, which impact their phonological development. Neural rhythmic speech studies have reported atypical delta phase in children with dyslexia, but beta band effects have not yet been studied. It is known that delta phase modulates the amplitude of the beta band response during rhythmic tasks via delta-beta phase-amplitude coupling (PAC). Accordingly, the atypical delta band effects reported for children with dyslexia may imply related atypical beta band effects. Here we analyse EEG data collected during a rhythmic speech paradigm from 51 children (21 typically-developing; 30 with dyslexia) who attended to a talking head repeating “ba” at 2Hz. Phase entrainment in the beta band, angular velocity in the beta band, power responses in the beta band and delta-beta PAC were assessed for each child and each group. Phase entrainment in the beta band was only significant for children without dyslexia. Children with dyslexia did not exhibit any phase consistency, and beta-band angular velocity was significantly faster compared to control children. Power in the beta band was significantly greater in the children with dyslexia. Delta-beta PAC was significant in both groups. The data are interpreted with respect to temporal sampling theory.

## 1. Introduction

There is an intimate developmental relationship between rhythmic skills and language development, and this relationship has been documented by behavioural studies in a range of languages (Fiveash et al., 2021, for recent review). Infants appear to begin the task of language-learning by utilising speech rhythm (Mehler et al., 1988), and linguistic analyses propose that languages can be divided into specific rhythm types (stress-timed, syllable-timed, moraic timing) which are already perceived as different by newborn infants (Nazzi et al., 1998). Indeed, computational modelling of infant-directed speech (IDS) shows that IDS has extra modulation energy in the delta band in the speech amplitude envelope compared to adult-directed speech, which serves perceptually to exaggerate rhythmic structure (Leong et al., 2017). Further, delta-band entrainment to rhythmic speech is significantly stronger than theta-band entrainment for infants aged 4, 7 and 11 months, and individual differences in delta-band cortical tracking at 11 months predict language outcomes at 24 months (Attaheri et al., 2022, 2023). The ubiquitous role of rhythm in language acquisition has led to the proposal that atypical rhythm skills in infancy and early childhood may be a core factor in the development of language disorders such as developmental dyslexia and developmental language disorder (DLD, Ladányi et al., 2020). Children with developmental dyslexia typically present with phonological difficulties relating to the sound structure of speech, while children with DLD present with grammatical impairments. At the sensory/neural level, this ‘atypical rhythm risk hypothesis’ is captured by Temporal Sampling (TS) theory (Goswami, 2011; 2015, 2022), which is focused on automatic processing of the speech amplitude envelope (AE).

TS theory suggests that difficulties in rhythm perception and production in infancy and childhood arise from atypical sensory/neural processing of the lower frequency portion (<10 Hz) of the AE, the slow-varying energy contour of speech which determines the perception of speech rhythm (Greenberg, 2006). Regarding the sensory aspects of TS theory, there are known to be different rates of amplitude modulation (AM) nested in the speech envelope centred on ∼2 Hz, ∼5 Hz and ∼20 Hz (Leong and Goswami, 2015), and research has shown that rates <10 Hz are those most critical for perceiving rhythm (Greenberg, 2003, 2006). Speech modelling of the AE (of IDS and nursery rhymes) has revealed that the *phase relations* between these different AM rates nested in the envelope (which approximately match the EEG bands of delta [0.5-4 Hz], theta [4-8 Hz], and beta/ low gamma [12-40 Hz] for child speech) provide systematic statistical clues to phonological units in speech (Leong et al., 2017; Leong and Goswami, 2015). Accordingly, difficulties in perceiving and encoding the different rates of AM nested in the speech AE and their phase relations could impair automatic statistical learning about the phonological structure of the speech signal for children with dyslexia. Computational analyses show that AM cycles at these 3 temporal rates (delta, theta, beta/low gamma) support the extraction of stressed vs unstressed syllable patterning (speech prosody), syllables, and onset-rimes respectively (‘acoustic-emergent phonology’, see Leong and Goswami, 2015). Children with dyslexia show impaired discrimination of amplitude rise times and AM across languages (Goswami, 2022, for recent review). The cognitive hallmark of developmental dyslexia is the “phonological core deficit” (Stanovich, 1988), defined by children’s reduced ability to identify and manipulate phonological units like syllables, rhymes and phonemes in oral tasks. By TS theory, this core phonological deficit arises in part from inefficient sensory discrimination of amplitude rise times and AMs.

Regarding the neural side of TS theory, MEG studies of children with dyslexia using story listening tasks suggest that the neural encoding impairments regarding speech information lie primarily in the encoding of low-frequency envelope information (Molinaro et al., 2016: Spanish; Mandke et al., 2022: English; Destoky et al., 2020; French). EEG studies assessing speech encoding mechanisms by using linear methods for measuring neural encoding such as temporal response functions (TRFs) suggest that children with dyslexia show atypical neural encoding of continuous speech in the delta band (Power et al., 2016; Di Liberto et al., 2018; Keshavarzi et al., 2022b). Potential differences between children with and without dyslexia in the neural phase of oscillation for delta, theta and alpha bands have been assessed by utilising rhythmic speech paradigms (such as repetition of the syllable ‘ba’ at 2 Hz; Power et al., 2012, 2013; Keshavarzi et al., 2022a). These latter EEG studies suggest that one core neural problem during speech processing relates to inaccurate phase synchronisation of the cortical networks that respond to delta-band AM information in the speech signal (Goswami, 2022, for a recent review). If the dyslexic brain is “out of phase” in its response to speech information in the delta band, then this might affect the accuracy of neural processing of the entire AM hierarchy nested in the speech signal, and thereby the extraction of phonology. This follows because neural responses to natural speech exhibit a mechanistic hierarchy governed by the delta band, in which phase relations and phase-amplitude relations between different frequency bands are utilised to capture the full complexity of the sensory information in speech (Gross et al., 2013; Ding and Simon, 2014; Doelling et al., 2014). For example, delta-beta PAC is thought to underpin temporal prediction accuracy during rhythmic processing via sensory-motor coupling (Arnal and Giraud, 2012; Arnal et al., 2015). Delta-beta PAC is already online in 4-month-old infants during speech listening (Attaheri et al., 2022), and is known to play a role in natural speech processing in the adult brain (Keitel et al., 2018). A prior study of children with dyslexia using a sentence listening task found intact delta-beta PAC (Power et al., 2016). Accordingly, atypical phase in the delta band might not necessarily be expected to attenuate delta-beta PAC, however this has yet to be investigated in rhythmic tasks.

Despite its potential importance for accurate rhythmic processing, the beta band has been relatively neglected in developmental language studies using EEG. However, recent longitudinal research in Mandarin Chinese based on resting-state EEG has shown that beta power increases over time between the ages of 7 and 11 years, and that the degree of increase in beta power from age 7 to 9 years predicts vocabulary at age 11 years (Meng et al., 2021). This relationship is significant even after controlling for cognitive ability, SES and home literacy environment (Meng et al., 2021). A role for the beta band in linguistic development may be consistent with adult data, as the magnitude of beta oscillatory responses in the adult brain has been associated with both speech comprehension and with predictive coding of speech (Gisladottir et al., 2018). Regarding beta band studies including children with dyslexia, a study of German children with and without developmental dyslexia found some group differences in beta-band power when the children were reading single words or pseudowords aloud (Klimesch et al., 2001). In particular, Klimesch and colleagues reported greater power in a narrow beta band range (12-14 Hz) in frontal, right central and centroparietal regions during pseudoword reading by children with dyslexia. However, the authors noted that the functional significance of this power increase was difficult to ascertain. Working in Italian, Spironelli et al. (2008) also reported greater beta-band power in children with dyslexia, both during a phonological task (judging whether two written words rhymed) and a semantic task (judging whether two written words were semantically related). Accordingly, developmental data suggest an involvement of beta power in various linguistic tasks delivered via print. More recently, Power et al. (2016) reported greater beta power in children with dyslexia compared to age-matched control children during a sentence listening task where no print was involved. In a word recognition task conducted with adults with dyslexia involving print (lexical decision task), Milne et al. (2003) recorded EEG while participants decided whether letter strings were words or not. They also reported higher beta power for dyslexic adults compared to non-dyslexic adults. Finally, in a recent EEG study with dyslexic adults using a beat perception task (participants listened to isochronous tones presented at a 2 Hz rate), Chang et al. (2021) reported that fluctuations in beta power differed between dyslexic and control adults. This was interpreted to reflect behavioural deficits in perceiving and tracking auditory rhythm in dyslexia, as would be predicted by TS theory.

Phase entrainment in the beta band has yet to be explored in children with dyslexia in a beat-based task. Accordingly, here we re-analyse the EEG data reported by Keshavarzi et al. (2022a), which investigated the delta, theta and alpha bands, this time focusing on the beta band. The EEG was recorded while children aged 9 years with and without dyslexia listened to the speech syllable “ba” presented rhythmically at a rate of 2 Hz. Given that the participating children with dyslexia had already shown less consistent phase entrainment in the delta band (Keshavarzi et al., 2022a) and given that delta phase couples with beta amplitude via delta-beta PAC (see also Arnal et al., 2014), group differences (in terms of phase entrainment, angular velocity, and beta power) in the beta band might be expected. We thus also computed delta-beta PAC, beta power and angular velocity in the beta band for our children. In the delta band analyses conducted with the same children, Keshavarzi et al. (2022a) reported a significantly greater angular velocity in children with dyslexia compared to control children over a time interval of –130 ms to 0 ms. This was taken to indicate that pre-stimulus delta phase was different in the two groups. If the phase of beta entrainment is atypical in the dyslexic brain, angular velocity in the beta band may also be expected to differ by group.

## 2. Method

### 2.1. Participants

Thirty children with developmental dyslexia (mean age = 110.7 months; SD = 5.6 months) and twenty-one typically developing children (mean age = 109.3 months; SD = 5.4 months) took part in the EEG study. The children were identified as dyslexic or typically-developing based on standardized reading, spelling and phonological awareness tests administered in 2018 (please see Keshavarzi et al., 2022a, for full details). They were assessed using the British Ability Scales standardized tests of reading and spelling (Elliott et al., 1996), the Test of Word Reading Efficiency word and nonword scales (TOWRE, Torgesen et al., 1999), and the rhyming subtest of the Phonological Assessment Battery (PhAB, Frederickson et al., 1997). Only children who scored at least one standard deviation below the test norm of 100 on at least two of the four reading and spelling measures and/or the phonology measure were included in the study. A summary of performance by group is shown as Table 1. Dyslexic children had no additional learning issues (e.g., dyspraxia, ADHD, autistic spectrum disorder, developmental language disorder) and were recruited through learning support teachers. They had a nonverbal IQ above 84, and English was their first language spoken at home. The absence of additional learning difficulties was assessed based on school and parental reports and our own testing impressions. Participants were attending state schools (equivalent to US public schools) situated in a range of towns and villages near a university town in the United Kingdom. Most families were Caucasian and of lower class or middle-class regarding income. All children received a short hearing screen using an audiometer. Sounds were presented in both the left and right ear at a range of frequencies (250, 500, 1000, 2000, 4000, 8000Hz), and all children were sensitive to sounds within the 20 dB HL range.

**Table 1.** Group characteristics expressed as mean and (S.D.) for children with dyslexia and age-matched control children on the reading, spelling and phonological measures.

|  | Dyslexic | Age-Matched Control |
| --- | --- | --- |
| N | 30 | 21 |
| Age (months) | 110.7 (5.6) | 109.3 (5.4) |
| BAS Reading SS | 81.0 (8.0) *** | 99.5 (6.2) |
| BAS Reading Age in months | 85.5 (11.0) *** | 106.5 (11.7) |
| BAS Spelling SS | 79.9 (7.5) *** | 97.1 (6.1) |
| TOWRE SWE SS | 79.5 (12.8) *** | 101.1 (7.7) |
| TOWRE PDE SS | 79.2 (10.9) *** | 98.0 (8.6) |
| PhAB Rhyme SS | 92.6 (11.7) *** | 102.4 (5.9) |

Note. \*\*\* $p < .001$ . BAS = British Ability Scales; SS = standardized score (mean = 100); TOWRE SWE = Test of Word Reading Efficiency Sight Word Efficiency Scale; TOWRE PDE = Phonic Decoding Efficiency Scale; PhAB = Phonological Assessment Battery.

### 2.2. Experimental paradigm and stimuli

The experimental paradigm and stimuli were exactly as described by Keshavarzi et al. (2022a). The stimuli comprised of rhythmic sequences of the syllable “ba” repeating 14 times at a rate of 2 Hz. One of the 14 “ba” syllables (at either position 9, 10 or 11) in each sequence was randomly out of time. Each child was presented with 90 trials, 15 of which were catch trials which were presented randomly and did not contain a lagged syllable. Each trial (except catch trials) consisted of three periods: the entrainment period (including syllables 1 – 8, 1 – 9 or 1–10 depending on the position of the lagged syllable), the violation period (including syllables 9 – 10, 10 – 11 or 11 – 12 depending on the position of the lagged syllable) and the return-to-isochrony period (including syllables 11 – 14, 12 – 14 or 13 – 14 depending on the position of the lagged syllable). Figure 1 illustrates an example of a trial with the lagged syllable at position 10. There was a fixed time interval of 500 ms between successive “ba” stimuli in the entrainment period. The extent to which the violator was out of the isochronous rhythm varied depending on how well the child responded in the task and followed a 3-down 1-up staircase procedure (Levitt, 1971). If a child correctly detected the violators in three successive trials, then the deviation from 500 ms stimulus-onset asynchrony decreased by 16.67 ms. If a violator was not detected, the deviation increased by 16.67 ms. There was a 3 second interval between successive trials.

**Figure 1.**
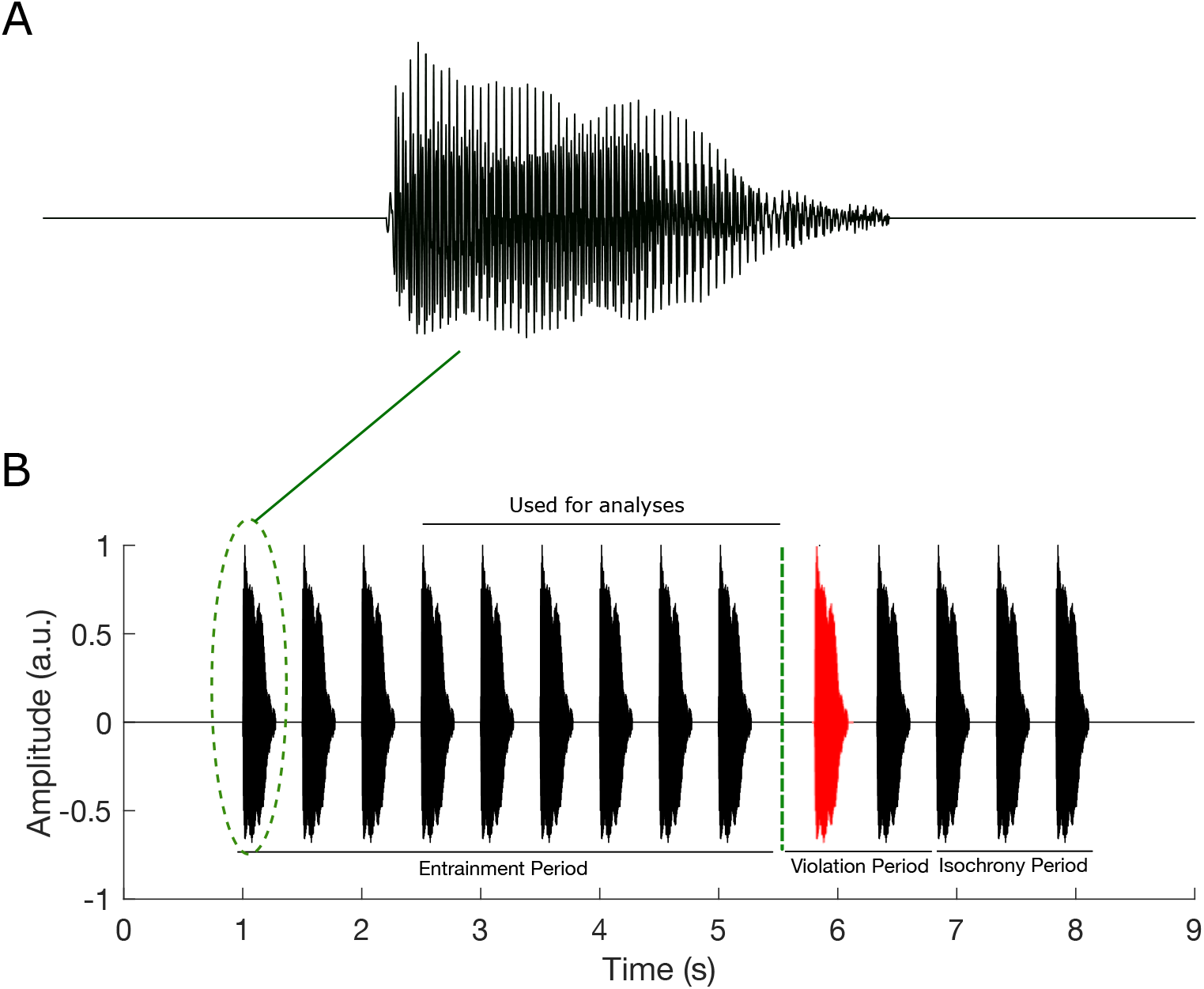
Experimental stimuli. Figure reproduced with permission from Keshavarzi et al., 2022a. (A) waveform of a single auditory “ba”, (B) a sequence of “ba” with the oddball as stimulus 10. The auditory stimuli consisted of the syllable “ba” repeated 14 times per trial at a rate of 2 Hz, with one syllable of stimuli 9-11 being out of the rhythm (here syllable 10, depicted in red). For this example (panel B), stimuli 4-9 of the entrainment period were used for the analysis.

### 2.3. EEG Data Acquisition

Participants were seated in a soundproof chamber. The auditory stimuli and visual speech information were presented to the participant and EEG data were simultaneously collected at 1-kHz sampling rate using a 128-channel EEG system (HydroCel Geodesic Sensor Net). Visual cues started 68 ms before the onset of the auditory stimulus “ba” as in natural speech. Children were instructed to focus on the lips of the talking face and to listen to the auditory stimuli. They also were asked to press a target key on the computer keyboard if one of the syllables was out of time. The EEG study (excluding the EEG set-up and preparation) took approximately 15 minutes.

### 2.5. EEG Data Pre-processing

The EEG pre-processing applied in this study is the same as the one described by Keshavarzi et al. (2022a). After considering the Cz channel as the reference, a zero phase FIR filter with a low cutoff (−6 dB) of 0.25 Hz and high cutoff (−6 dB) of 48.25 Hz (EEGLab Toolbox; Delorme and Makeig, 2004) was used to band-passed filter data at frequency range of 0.5 – 48 Hz. We detected extremely noisy channels using the spectrogram, kurtosis and probability methods provided by EEGLAB Toolbox. If a channel was 3 S.D. away from the average, it was rejected and interpolated using spherical interpolation (EEGLab Toolbox). We applied the independent component analysis, provided in EEGLab Toolbox, to the data for each participant. The obtained independent components were then assessed carefully to remove artefactual components such as eye movements. Participant head movements during the EEG recording were monitored using a camera installed in the EEG room and the times corresponding to head movements were manually recorded by the experimenter. This information was then used to ensure that trials corrupted by head movements had been cleaned by the pre-processing steps. The EEG data were downsampled to 100 Hz and band-pass filtered to extract delta (0.5 – 4 Hz) and beta (15 – 25 Hz) frequency bands. The data were then epoched into individual trials, in the time range of 500 ms before the onset of the first stimulus and 4.5 sec after that. Some trials were excluded from the analyses because of technical issues such as stimulus information not being marked in the EEG file. The average number of trials utilised for the analyses for control and dyslexic groups were 77.61 (S.D. = 20.01) and 78.13 (S.D. = 21.67), respectively. We obtained the instantaneous phase separately for each EEG channel (filtered at the beta band) using two steps: (1) Computing the analytic representation of the input signal through the Hilbert transform; (2) Computing the phase of each sample (which is a complex value) of the analytic signal. Note that the focus of the analyses in this study was only on the entrainment period. To ensure that rhythmicity had been established and to maximise data, we excluded the first two “ba” stimuli for trials with a violation occurring at position 9 (hence our analyses used stimuli 3 – 8) and the first three “ba” stimuli for trials where the violation occurred at position either 10 or 11 (hence our analyses used stimuli 4 – 9). This resulted in a maximum of 540 “ba” stimuli for each participant.

### 2.7. Computation of phase entrainment

To assess the phase entrainment for each group in the beta band, the following six steps were performed (as previously used by Keshavarzi et al. (2022a) for investigating delta, theta and alpha):

- Step 1. Calculating instantaneous phases of all 128 EEG channels at the times corresponding to the onsets of the 6 “ba” stimuli that were used for the analyses for each of the 90 trials.
- Step 2. Calculating the mean phase for each of the 90 trials by averaging across the phase observations obtained for 128 EEG channels and for the 6 “ba” stimuli in step 1. This results in a single-phase value for each trial.
- Step 3. Deriving a single unit vector (whose angle is determined by the phase value obtained in step 2) in the vector space for each trial.
- Step 4. Calculating the mean vector for each child by averaging across the unit vectors obtained in step 3. This results in a single vector for each child, subsequently we refer to this as the *child resultant vector*. The length of the *child resultant vector* is a value between 0 and 1 and its angle, called *child preferred phase*, is between 0 and 2π. The length of the *child resultant vector* can be used as a criterion to assess the strength of phase consistency across different trials for each individual participant.
- Step 5. A single unit vector (whose angle is determined by the *child preferred phase* obtained in step 4) is considered in the vector space for each child.
- Step 6. The mean vector for each group is computed by averaging across the unit vectors obtained in step 5. This produces a single vector, called the *group resultant vector*, for each group. The length of the *group resultant vector* is a value between 0 and 1 and its angle, called *group preferred phase*, is between 0 and 2π. The length of the *group resultant vector* can be used as a criterion to assess the strength of phase consistency across different children by group.

### 2.8. Computation of phase-amplitude coupling

Phase amplitude coupling (PAC) refers to a type of cross-frequency coupling in which the amplitude of the signal at a high-frequency band is modulated by the phase of low-frequency oscillations. In this study, we quantified the strength of this modulation using the modulation index (*MI*; Tort et al. 2008; Hülsemann, 2019):

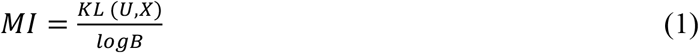

where *B* (=18) is the number of bins, *U* refers to the uniform distribution, *X* is the distribution of the data, and *KL (U,X)* is Kullback–Leibler distance, a measure of the disparity of two distributions (Hülsemann, 2019), and is calculated by:

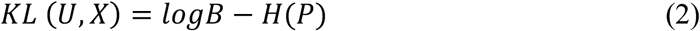

where *H*(•)is the Shannon entropy and *P* is the vector of normalized averaged amplitudes per phase bin which is calculated as:

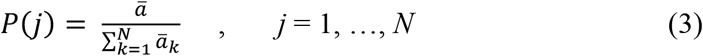

where *ā* is the average amplitude of each bin and *k* refers to running index for the bins. Note that *P* is a vector with *N* elements.

## 3. Results

### 3.1. Phase entrainment in the beta band within each group

To check the consistency of phase entrainment across the different children in each group, the Rayleigh test of uniformity was applied to the *child preferred phases* (angles of *child resultant vectors*, see step 4 in subsection 2.7) separately for each group. The distribution of phase at the onsets of the “ba” stimuli was significantly different from the uniform distribution for the control children (Rayleigh test, *z* = 4.13, *p* = 0.01; see Figure 2A), indicating consistent phase entrainment. For children with dyslexia, however, the distribution of phase at the onsets of the “ba” stimuli did not differ from the uniform distribution (Rayleigh test, *z* = 0.27, *p* = 0.76; see Figure 2B), suggesting that consistent phase entrainment was not present in the beta band.

**Figure 2.**
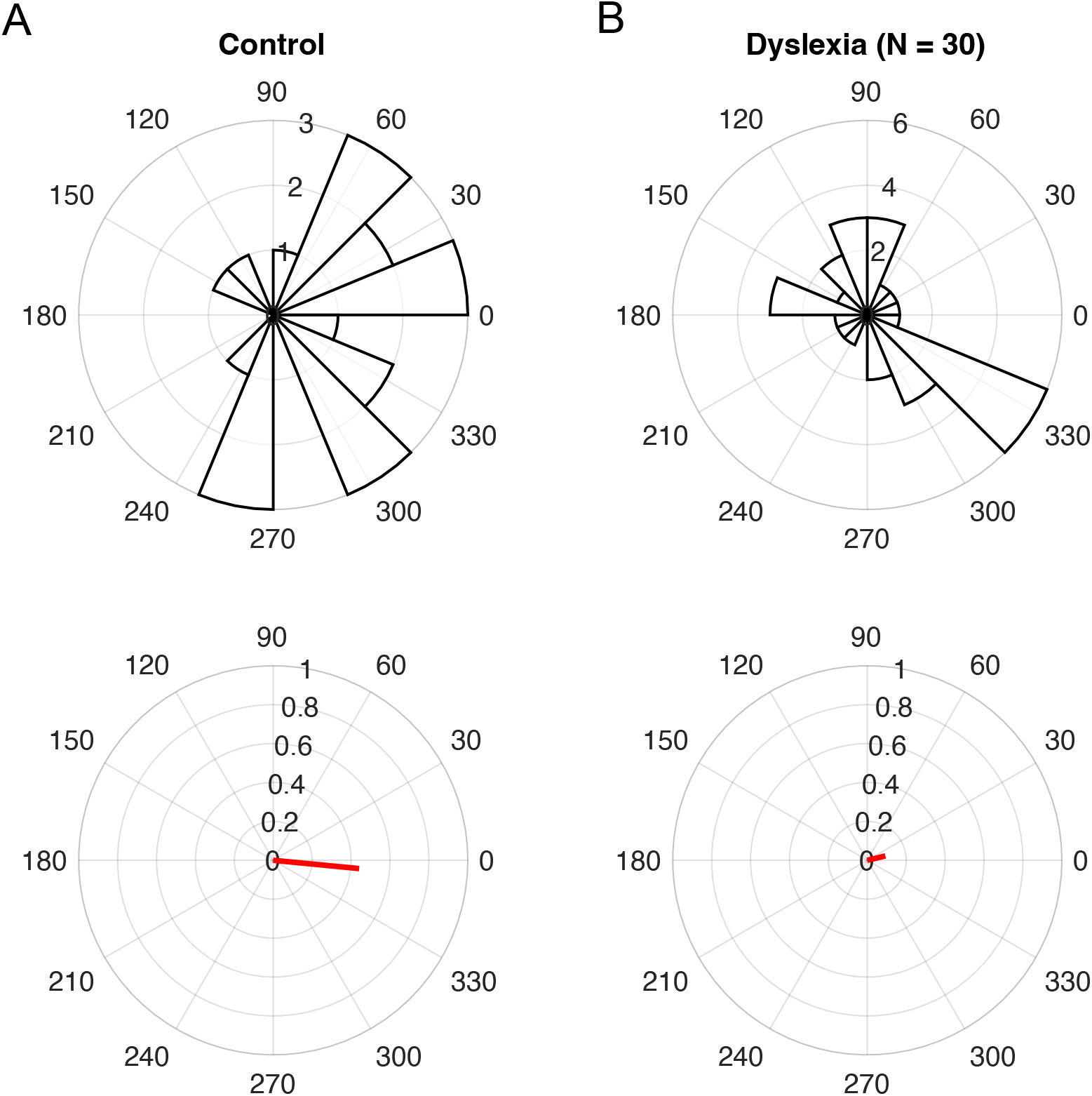
Phase distribution across groups and group resultant vectors (red lines). There is significant phase entrainment across different children in the control group (panel A) but not in the dyslexic group (Panel B). The length of the group resultant vector can be used as a criterion to assess the strength of phase entrainment.

### 3.2. Comparing angular velocity between groups

To further assess the behaviour of preferred phase in the beta band for each group, we computed angular velocity. Angular velocity is the rate of phase changes across time, providing an index of pre-stimulus differences in phase just before the occurrence of the next “ba” syllable. *Group angular velocity* was computed separately for each group in the entrainment period over the time interval of –500 ms to 500 ms (relative to the occurrence of a stimulus, see Figure 3). Following Keshavarzi et al.’s (2022a) report of a significant group difference in terms of pre-stimulus angular velocity, we used two-sample t-tests to check for a group difference in terms of angular velocity over different time intervals. The results showed that there was a significant difference in angular velocity between the two groups over the time interval of –350 ms to –90 ms (controls: 34.2π rad/s; children with dyslexia: 37.6π rad/s; two-sample t-test, *p* = 0.008). Accordingly, in the prestimulus period, the rate of beta phase changes differs significantly between the two groups. These changes were faster in the children with dyslexia.

**Figure 3.**
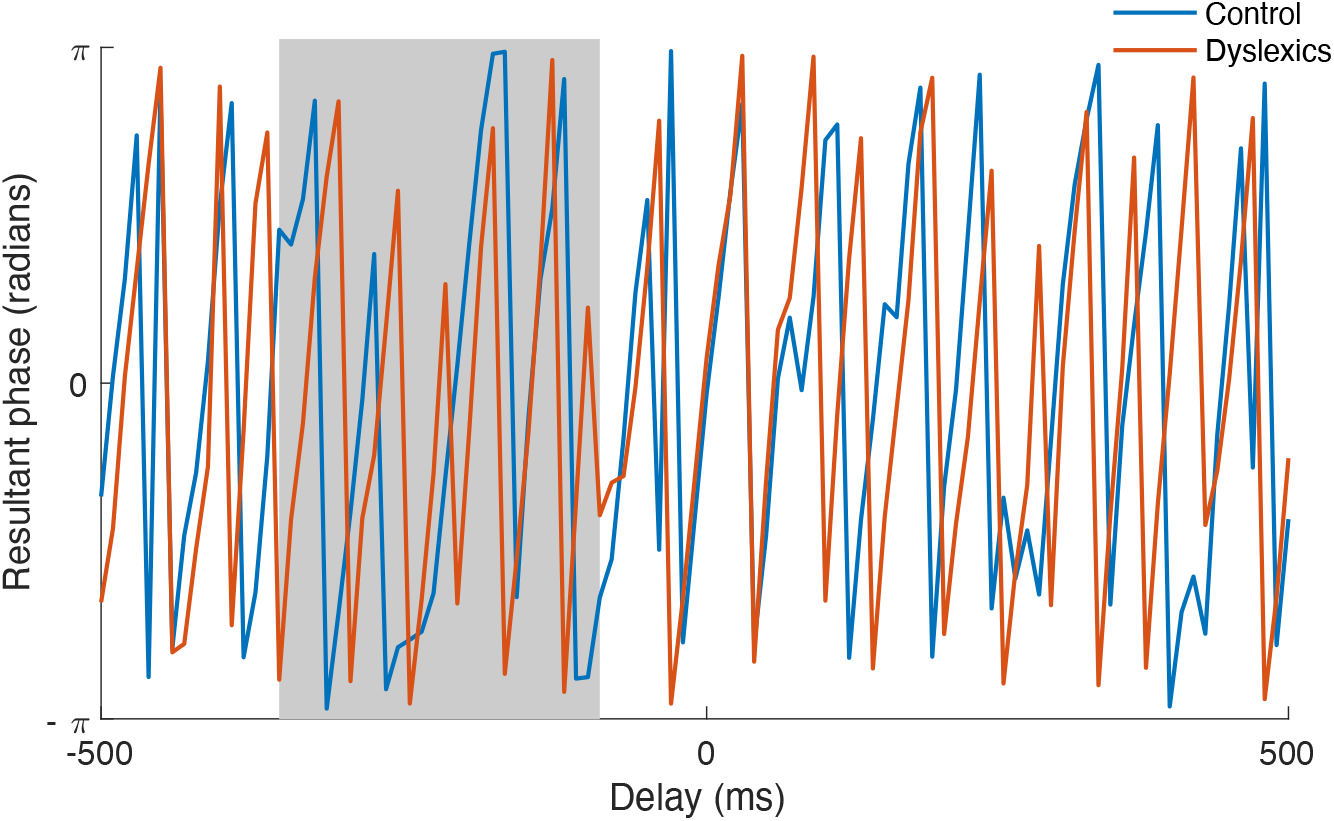
The angular velocity (radians) versus the delay (ms) relative to “ba” stimuli onsets. The blue and red curves are for control and dyslexic groups, respectively.

### 3.3. Beta band power over the entrainment period (3 seconds)

To investigate the beta band power of neural response over the entrainment period used for analysis (3 seconds), we calculated the power of EEG responses for each child and each group separately. This was conducted in four steps: (1) Calculating the power of each EEG channel separately for each trial; (2) Calculating the power for each trial by averaging across power values obtained (in step 1) for channels; (3) Calculating the power for each child by averaging across power values obtained (in step 2) for all trials of that child; (4) Calculating the power for each group by averaging across power values obtained (in step 2) for all children in that group. Figure 4 demonstrates the beta band power for each group. To compare average beta-band power for children in the control group with those in the dyslexic group, we applied the Wilcoxon rank sum test after removing outliers. Data points were treated as outliers if the corresponding power value was more than a 1.5 interquartile range above the upper quartile or below the lower quartile of the population data in that group. There were 4 outliers in the dyslexic group and 3 outliers in the control group. The results showed that beta band power obtained for the children with dyslexia was significantly greater than that of control children (Wilcoxon rank sum test, *z* = -2.42, *p* = 0.015).

**Figure 4.**
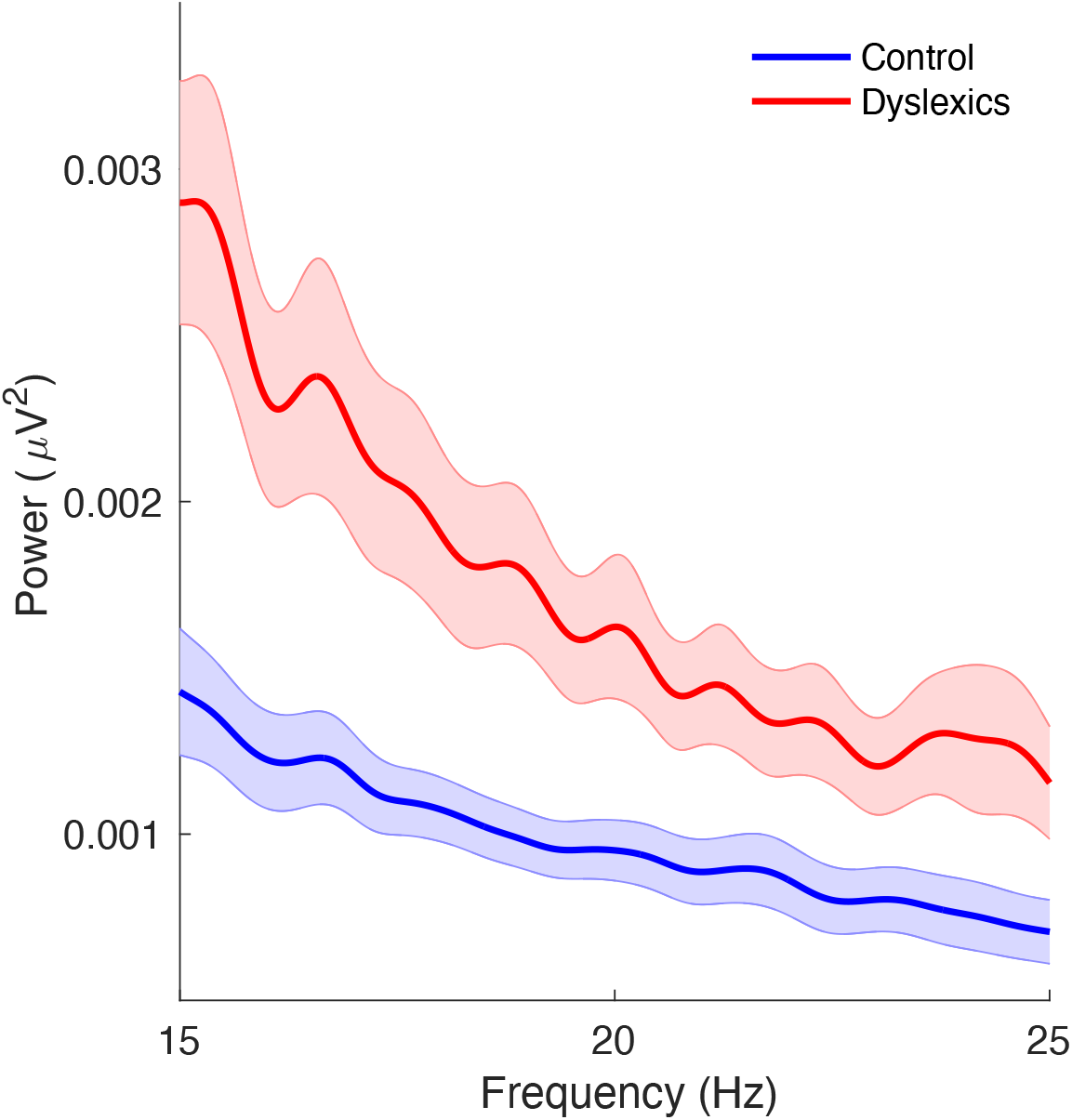
Beta-band power. The blue and red curves denote the power in the beta band versus frequency for control and dyslexic groups, respectively. The shaded areas demonstrate the standard error of the mean.

### 3.4. Delta-beta phase-amplitude coupling (PAC)

To study delta-beta PAC, the *MI* was computed separately for all trials of each child. The mean *MI* for a given child was then calculated by averaging across the *MI* scores obtained from the trials completed by that child. Figure 5 shows the mean *MI* scores for children in each group (control versus dyslexic). To compare *MI* between the two groups, a two-sample t-test was applied. No significant difference in PAC between the two groups was observed (Two-sample t-test, *p* = 0.46).

**Figure 5.**
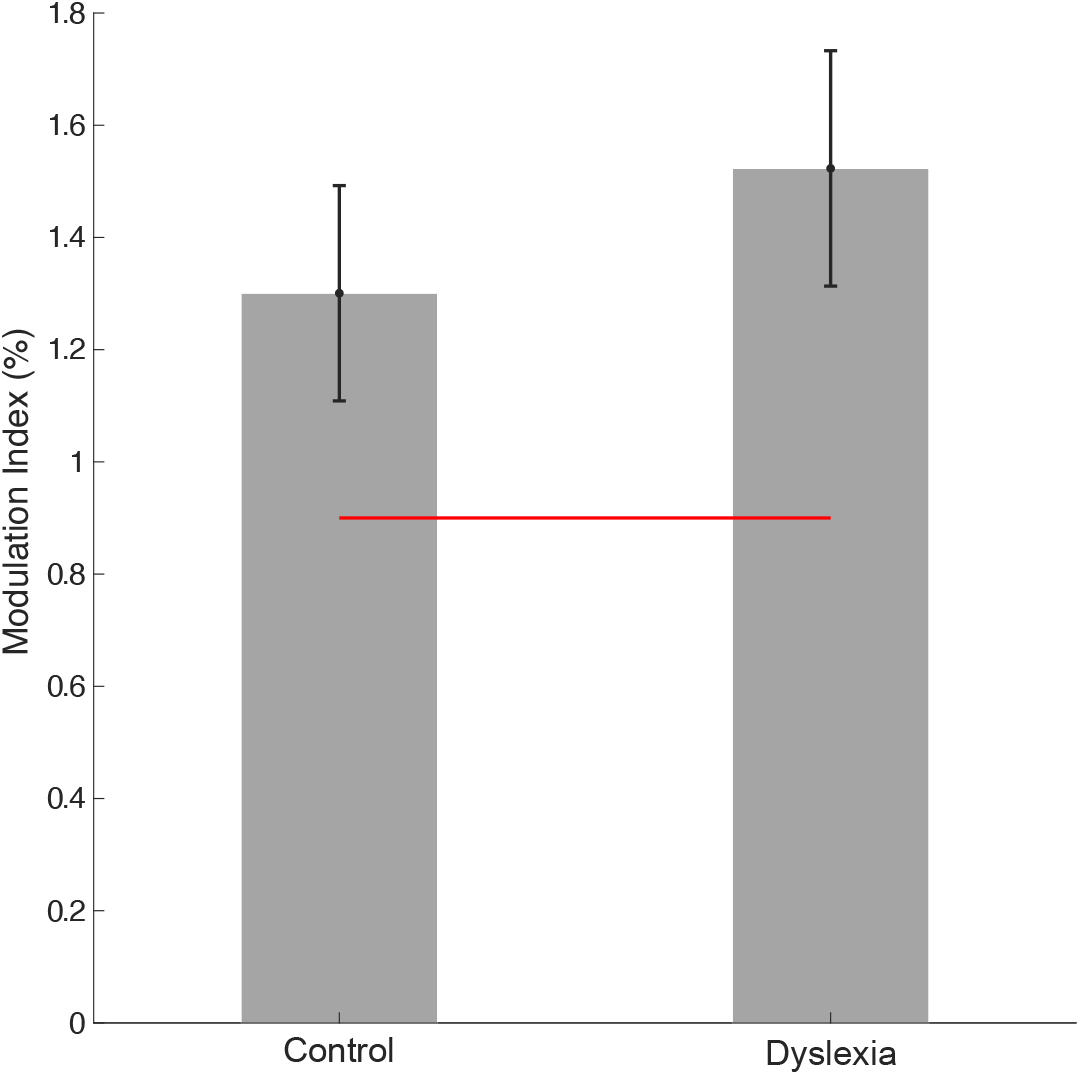
Modulation index as a measure of the strength of delta-beta phase-amplitude coupling for control and dyslexic group. The horizontal red line denotes the chance level.

To assess whether the *MI* values obtained for dyslexic and control children were above chance, null models were computed. To this end, the EEG data corresponding to all 90 trials was first replaced by red noise (Hülsemann, 2019). We then calculated the *MI* separately for each trial using the procedure described in Subsection 2.8. We next computed the mean *MI* value by averaging across *MI* scores obtained for all 90 trials, resulting in one single score. This procedure was performed 1000 times, yielding 1000 scores. Chance level was obtained by averaging across these 1000 scores. To statistically compare *MI* scores obtained for children in the control group and in the dyslexic group with chance, we applied Wilcoxon rank sum tests separately. The results showed that *MI* scores obtained for both children with dyslexia and control children were statistically greater than chance (Wilcoxon rank sum test; control: *z* = 5.61, *p* = 2×10^−8^; dyslexia: *z* = 8.1, *p* = 5×10^−16^). Accordingly, significant delta-beta PAC was present in both groups.

## 4. Discussion

Here we set out to explore potential group differences in beta phase, beta power, angular velocity and delta-beta PAC between children with and without dyslexia in response to rhythmic speech. Given that prior studies with rhythmic speech as input have established atypical phase entrainment in the delta band for children with dyslexia (Power et al., 2013; Keshavarzi et al., 2022a), and given that delta phase modulates beta amplitude (through delta-beta PAC) during speech processing for children (Power et al., 2016), it was expected that some atypical effects in the beta band may also be present in those participants with developmental dyslexia. The data showed that while significant phase entrainment in the beta band was present for the control children, the children with dyslexia did not show significant phase entrainment in the beta band (Figure 2). Accordingly, by the age of 9 years, the dyslexic brain has not yet established phase consistency in the beta band when responding to rhythmic speech at 2 Hz. However, group angular velocity did show a significant difference between groups. Group angular velocity over the time interval of – 500 ms to 500 ms (relative to the occurrence of a stimulus, see Figure 3) was investigated separately for each group. There was a significant difference in angular velocity between the two groups over the time interval of –350 ms to –90 ms (see Figure 3), indicative of faster changes in beta phase in the children with dyslexia. Accordingly, beta phase in response to rhythmic input does appear to be atypical in the dyslexic brain.

We also explored delta-beta PAC. PAC mechanisms are assumed to support the integration (coordination) of processes in the brain (Mariscal et al., 2021), and delta-beta PAC has been particularly studied with regard to temporal processing. Concerning rhythmic processing, Arnal et al. (2014) have shown that oscillations in the delta and beta bands are instrumental in temporal prediction in the adult brain, aligning with upcoming targets and supporting accurate behavioural responding. Concerning speech processing, Pefkou et al. (2017) reported a role for endogenous beta oscillations in speech comprehension by adults using a time-compressed speech paradigm. Beta oscillations in the motor cortex contribute to planning movements, and Morillon et al. (2019) showed functional delta-beta coupling during planned motor acts, which they argued may provide a contextual temporal framework for processing slower linguistic information. In the current rhythmic speech study, delta-beta PAC was found to be intact for the 9-year-old children with dyslexia (see Figure 5). A similar result in a sentence listening paradigm was previously reported for 14-year-old children with developmental dyslexia by Power et al. (2016). However, this finding does not necessarily suggest that motor-based temporal prediction mechanisms in the dyslexic brain are functioning efficiently. Indeed, it is known that children with dyslexia have difficulty in tapping in time with a beat (Thomson and Goswami, 2008; Colling et al., 2017). In a reaction time task based on rhythmic tones, adults with dyslexia do not show the same relation between faster reaction times and the rising phase of the delta oscillation that is shown by non-dyslexic adults, despite being matched for overall performance (Soltész et al., 2013). In adult studies, the entrained delta-band phase-course has been thought to provide an internal “chronograph”, accurately tracking elapsing time via the accumulation of phase information (Arnal and Kleinschmidt, 2017). In particular, delta phase is known from adult studies to be important in detecting audio-visual temporal asynchronies when prosodic information is manipulated (Biau et al., 2022). Children with dyslexia have documented prosodic difficulties and show atypical delta phase (Goswami, 2022, for review), yet such prosodic temporal and motor paradigms have not yet been utilised with children with dyslexia. Intriguingly, Biau et al. (2022) noted decreased delta-beta coupling in left motor cortex when their adult participants could not accurately map visual and auditory prosodies. It is thus possible that more naturalistic speech tasks than the rhythmic speech task employed here could reveal some differences in delta-beta PAC in dyslexia.

Meanwhile, prior studies with participants with dyslexia have documented greater beta power for both children with dyslexia (Klimesch et al., 2001; Spironelli et al., 2008) and adults with dyslexia (Milne et al., 2003) during reading-related tasks. In the current speech listening study, beta power in the dyslexic brain was significantly greater than beta power for the control participants, replicating a finding previously reported by Power et al. (2016) for sentence listening (see Figure 4). Beta-band power is known to be related to visual attention (Gola et al., 2013). Accordingly, in the current audio-visual speech paradigm, greater beta band power in dyslexia may serve as a compensatory mechanism for the impaired neural sampling of auditory information. This could be investigated directly in future studies. Finally, although our task (listening to syllables presented at a 2 Hz rate) is similar to the rhythmic task used by Chang et al. (2021) with dyslexic adults (listening to tones presented at a 2 Hz rate), the main focus of our study was on the phase entrainment of the neural responses in the beta band, and not on the phase of beta band power fluctuation. As the children with dyslexia tested here did not show any phase consistency in the beta band, we could not analyse the phase of beta band power fluctuations.

The current study has a number of limitations. Due to the Covid-19 Pandemic, the number of control children was fewer than the number of children with dyslexia. Secondly, the paradigm did not involve naturalistic continuous speech. Future studies with rhythmic tasks, ideally using naturalistic speech paradigms such as the prosodic tasks devised by Biau et al. (2022), are required in order to deepen our understanding of the potential role of beta oscillations in linguistic processing for participants with dyslexia.

## Ethics approval statement

All participants and their parents gave informed consent for the EEG study in accordance with the Declaration of Helsinki, and the study was reviewed by the Psychology Research Ethics Committee of the University of Cambridge.

## Data availability

Data will be made available on request.

## Acknowledgement

The authors would like to thank Barbara Tillmann for first suggesting that we investigate beta phase, and all the children, families and schools involved in the study.

## References

Arnal, L.H., Doelling, K.B., Poeppel, D., 2015. Delta–beta coupled oscillations underlie temporal prediction accuracy. Cerebral Cortex, 25(9), 3077–3085. https://doi.org/10.1093/cercor/bhu103.

Arnal, L.H., Giraud, A.L., 2012. Cortical oscillations and sensory predictions. Trends in cognitive sciences, 16(7), 390–398. https://doi.org/10.1016/j.tics.2012.05.003.

Arnal, L.H., Kleinschmidt, A.K., 2017. Entrained delta oscillations reflect the subjective tracking of time. Communicative & integrative biology, 10(5-6), 3077–3085. https://doi.org/10.1080/19420889.2017.1349583.

Attaheri, A., Choisdealbha, Á.N., Di Liberto, G.M., Rocha, S., Brusini, P., Mead, N.,, Goswami, U., 2022. Delta-and theta-band cortical tracking and phase-amplitude coupling to sung speech by infants. NeuroImage, 247, 118698. https://doi.org/10.1016/j.neuroimage.2021.118698.

Attaheri, A., Ní Choisdealbha, Á., Rocha, S., Brusini, P., Di Liberto, G.M., Mead, N.,, Goswami, U., 2022. Infant low-frequency EEG cortical power, cortical tracking and phase-amplitude coupling predicts language a year later. bioRxiv. https://doi.org/10.1101/2022.11.02.514963.

Biau, E., Schultz, B.G., Gunter, T.C., Kotz, S.A., 2022. Left motor δ oscillations reflect asynchrony detection in multisensory speech perception. Journal of Neuroscience, 42(11), 2313–2326.https://doi.org/10.1523/JNEUROSCI.2965-20.2022.

Chang, A., Bedoin, N., Canette, L.H., Nozaradan, S., Thompson, D., Corneyllie, A.,, Trainor, L.J., 2021. Atypical beta power fluctuation while listening to an isochronous sequence in dyslexia. Clinical Neurophysiology, 132(10), 2384–2390. https://doi.org/10.1016/j.clinph.2021.05.037.

Colling, L.J., Noble, H.L., Goswami, U., 2017. Neural entrainment and sensorimotor synchronization to the beat in children with developmental dyslexia: An EEG study. Neuroscience, 11, 360. https://doi.org/10.3389/fnins.2017.00360.

Delorme, A., Makeig, S., 2004. EEGLAB: an open source toolbox for analysis of single-trial EEG dynamics including independent component analysis. Journal of neuroscience methods, 134(1), 9–21. https://doi.org/10.1016/j.jneumeth.2003.10.009.

Destoky, F., Bertels, J., Niesen, M., Wens, V., Vander Ghinst, M., Leybaert, J.,, Bourguignon, M., 2020. Cortical tracking of speech in noise accounts for reading strategies in children. PLoS biology, 18(8), e3000840. https://doi.org/10.1371/journal.pbio.3000840.

Di Liberto, G.M., Peter, V., Kalashnikova, M., Goswami, U., Burnham, D., Lalor, E.C., 2018. Atypical cortical entrainment to speech in the right hemisphere underpins phonemic deficits in dyslexia. NeuroImage, 175, 70–79. https://doi.org/10.1016/j.neuroimage.2018.03.072.

Ding, N., Simon, J.Z., 2014. Cortical entrainment to continuous speech: functional roles and interpretations. Frontiers in human neuroscience, 8, 311. https://doi.org/10.3389/fnhum.2014.00311.

Doelling, K.B., Arnal, L.H., Ghitza, O., Poeppel, D., 2014. Acoustic landmarks drive delta–theta oscillations to enable speech comprehension by facilitating perceptual parsing. NeuroImage, 85, 761–768. https://doi.org/10.1016/j.neuroimage.2013.06.035.

Elliott, C.D., Smith, P., McCullogh, K., 1996. British Ability Scales (2nd ed.). Windsor, UK: NFER-Nelson.

Fiveash, A., Bedoin, N., Gordon, R.L., Tillmann, B., 2021. Processing rhythm in speech and music: Shared mechanisms and implications for developmental speech and language disorders. Neuropsychology, 35(8), 771. https://doi.org/10.1037/neu0000766.

Frederickson, N., Frith, U., Reason, R., 1997. Phonological assessment battery (PhAB): Manual and test materials. NFER-Nelson.

Gisladottir, R.S., Bögels, S., Levinson, S.C., 2018. Oscillatory brain responses reflect anticipation during comprehension of speech acts in spoken dialog. Frontiers in human neuroscience, 12, 34. https://doi.org/10.3389/fnhum.2018.00034.

Gola, M., Magnuski, M., Szumska, I., Wróbel, A., 2013. EEG beta band activity is related to attention and attentional deficits in the visual performance of elderly subjects. International Journal of Psychophysiology, 89(3), 334–341. https://doi.org/10.1016/j.ijpsycho.2013.05.007.

Goswami, U., 2011. A temporal sampling framework for developmental dyslexia. Trends in cognitive sciences, 15(1), 3–10. https://doi.org/10.1016/j.tics.2010.10.001.

Goswami, U., 2015. Sensory theories of developmental dyslexia: three challenges for research. Nature Reviews Neuroscience, 16(1), 43–54. https://doi.org/10.1038/nrn3836.

Goswami, U., 2022. Language acquisition and speech rhythm patterns: an auditory neuroscience perspective. Royal Society Open Science, 9(7), 211855. https://doi.org/10.1098/rsos.211855.

Greenberg, S., Carvey, H., Hitchcock, L., Chang, S., 2003. Temporal properties of spontaneous speech—a syllable-centric perspective. Journal of Phonetics, 31(3-4), 465–485. https://doi.org/10.1016/j.wocn.2003.09.005.

Gross, J., Hoogenboom, N., Thut, G., Schyns, P., Panzeri, S., Belin, P., Garrod, S., 2013. Speech rhythms and multiplexed oscillatory sensory coding in the human brain. PLoS biology, 11(12), e1001752. https://doi.org/10.1371/journal.pbio.1001752.

Hülsemann, M.J., Naumann, E., Rasch, B., 2019. Quantification of phase-amplitude coupling in neuronal oscillations: comparison of phase-locking value, mean vector length, modulation index, and generalized-linear-modeling-cross-frequency-coupling. Frontiers in neuroscience, 13, 573. https://doi.org/10.3389/fnins.2019.00573.

Keitel, A., Gross, J., Kayser, C., 2018. Perceptually relevant speech tracking in auditory and motor cortex reflects distinct linguistic features. PLoS biology, 16(3), e2004473. https://doi.org/10.1371/journal.pbio.2004473.

Keshavarzi, M., Mandke, K., Macfarlane, A., Parvez, L., Gabrielczyk, F., Wilson, A., Goswami, U., 2022a. Atypical delta-band phase consistency and atypical preferred phase in children with dyslexia during neural entrainment to rhythmic audio-visual speech. NeuroImage: Clinical, 35, 103054. https://doi.org/10.1016/j.nicl.2022.103054.

Keshavarzi, M., Mandke, K., Macfarlane, A., Parvez, L., Gabrielczyk, F., Wilson, A., …, Goswami, U., 2022b. Decoding of speech information using EEG in children with dyslexia: Less accurate low-frequency representations of speech, not “Noisy” representations. Brain and Language, 235, 105198. https://doi.org/10.1016/j.bandl.2022.105198.

Klimesch, W., Doppelmayr, M., Wimmer, H., Gruber, W., Röhm, D., Schwaiger, J., Hutzler, F., 2001. Alpha and beta band power changes in normal and dyslexic children. Clinical Neurophysiology, 112(7), 1186–1195. https://doi.org/10.1016/s1388-2457(01)00543-0.

Ladányi, E., Persici, V., Fiveash, A., Tillmann, B., Gordon, R.L., 2020. Is atypical rhythm a risk factor for developmental speech and language disorders?. Wiley Interdisciplinary Reviews: Cognitive Science, 11(5), e1528. https://doi.org/10.1002/wcs.1528.

Levitt, H.C.C.H., 1971. Transformed up-down methods in psychoacoustics. The Journal of the Acoustical society of America, 49(2B), 467–477. https://doi.org/10.1121/1.1912375.

Mandke, K., Flanagan, S., Macfarlane, A., Gabrielczyk, F., Wilson, A., Gross, J., Goswami, U., 2022. Neural sampling of the speech signal at different timescales by children with dyslexia. NeuroImage, 253, 119077. https://doi.org/10.1016/j.neuroimage.2022.119077.

Meng, X., Sun, C., Du, B., Liu, L., Zhang, Y., Dong, Q., …, Nan, Y., 2022. The development of brain rhythms at rest and its impact on vocabulary acquisition. Developmental science, 25(2), e13157. https://doi.org/10.1111/desc.13157.

Milne, R.D., Hamm, J.P., Kirk, I.J., Corballis, M.C., 2003. Anterior–posterior beta asymmetries in dyslexia during lexical decisions. Brain and Language, 84(3), 309–317. https://doi.org/10.1016/s0093-934x(02)00557-6.

Leong, V., Goswami, U., 2015. Acoustic-emergent phonology in the amplitude envelope of childdirected speech. PloS One, 10(12), e0144411. https://doi.org/10.1371/journal.pone.0144411.

Leong, V., Kalashnikova, M., Burnham, D., Goswami, U., 2017. The temporal modulation structure of infant-directed speech. Open Mind, 1(2), 78–90. https://doi.org/10.1162/OPMI_a_00008.

Mariscal, M.G., Levin, A.R., Gabard-Durnam, L.J., Xie, W., Tager-Flusberg, H., Nelson, C.A., 2021. EEG phase-amplitude coupling strength and phase preference: association with age over the first three years after birth. Eneuro, 8(3). https://doi.org/10.1523/eneuro.0264-20.2021.

Mehler, J., Jusczyk, P., Lambertz, G., Halsted, N., Bertoncini, J., Amiel-Tison, C., 1988. A precursor of language acquisition in young infants. Cognition, 29(2), 143–178. https://doi.org/10.1016/0010-0277(88)90035-2.

Molinaro, N., Lizarazu, M., Lallier, M., Bourguignon, M., Carreiras, M., 2016. Out-of-synchrony speech entrainment in developmental dyslexia. Human brain mapping, 37(8), 2767–2783. https://doi.org/10.1002/hbm.23206.

Morillon, B., Arnal, L.H., Schroeder, C.E., Keitel, A., 2019. Prominence of delta oscillatory rhythms in the motor cortex and their relevance for auditory and speech perception. Neuroscience & Biobehavioral Reviews, 107, 136–142. https://doi.org/10.1016/j.neubiorev.2019.09.012.

Nazzi, T., Bertoncini, J., Mehler, J., 1998. Language discrimination by newborns: toward an understanding of the role of rhythm. Journal of Experimental Psychology: Human perception and performance, 24(3), 756. https://doi.org/10.1037//0096-1523.24.3.756.

Pefkou, M., Arnal, L.H., Fontolan, L., Giraud, A.L., 2017. θ-Band and β-band neural activity reflects independent syllable tracking and comprehension of time-compressed speech. Journal of Neuroscience, 37(33), 7930–7938. https://doi.org/10.1523/jneurosci.2882-16.2017.

Power, A.J., Colling, L.J., Mead, N., Barnes, L., Goswami, U., 2016. Neural encoding of the speech envelope by children with developmental dyslexia. Brain and Language, 160, 1–10. https://doi.org/10.1016/j.bandl.2016.06.006.

Power, A.J., Mead, N., Barnes, L., Goswami, U., 2012. Neural entrainment to rhythmically presented auditory, visual, and audio-visual speech in children. Frontiers in Psychology, 3, 216. https://doi.org/10.3389/fpsyg.2012.00216.

Power, A.J., Mead, N., Barnes, L., Goswami, U., 2013. Neural entrainment to rhythmic speech in children with developmental dyslexia. Frontiers in human neuroscience, 7, 777. https://doi.org/10.3389/fnhum.2013.00777.

Soltész, F., Szűcs, D., Leong, V., White, S., Goswami, U., 2013. Differential entrainment of neuroelectric delta oscillations in developmental dyslexia. PloS One, 8(10), Article e76608. https://doi.org/10.1371/journal.pone.0076608.

Spironelli, C., Penolazzi, B., Angrilli, A., 2008. Dysfunctional hemispheric asymmetry of theta and beta EEG activity during linguistic tasks in developmental dyslexia. Biological psychology, 77(2), 123–131. https://doi.org/10.1016/j.biopsycho.2007.09.009.

Thomson, J.M., Goswami, U., 2008. Rhythmic processing in children with developmental dyslexia: Auditory and motor rhythms link to reading and spelling. Journal of Physiology-Paris, 102, 120–129. https://doi.org/10.1016/j.biopsycho.2007.09.009.

Torgesen, J., Wagner, R.K., Rashotte, C., 1999. Test of Word Reading Efficiency (TOWRE). Austin, TX: Pro-Ed.

Tort, A.B., Kramer, M.A., Thorn, C., Gibson, D.J., Kubota, Y., Graybiel, A.M., Kopell, N.J., 2008. Dynamic cross-frequency couplings of local field potential oscillations in rat striatum and hippocampus during performance of a T-maze task. Proceedings of the National Academy of Sciences, 105(51), 20517–20522. https://doi.org/10.1073/pnas.0810524105.

